# Untargeted Metabolomic Profiling Identifies Disease-specific Profiles in Food Allergy

**DOI:** 10.1101/657866

**Authors:** E Crestani, Hani Harb, Louis Marie Charbonnier, J Leirer, A Motsinger-Reif, Rima Rachid, W Phipatanakul, R Kaddurah-Daouk, T.A. Chatila

## Abstract

**Background:** Food allergy (FA) affects an increasing proportion of children for reasons that remain obscure. Identification of pathogenic mechanisms involved in FA using untargeted metabolomic approaches may provide much needed diagnostic and prognostic disease biomarkers and improved treatment options.

**Methods:** Mass spectrometry-based untargeted metabolomic profiling was performed on serum samples of children with either FA alone, asthma alone or both FA and asthma as well as healthy pediatric controls.

**Results:** FA subjects exhibited a disease-specific metabolomic signature as compared to both control subjects and asthmatics. In particular, FA was uniquely associated with a marked decrease in sphingolipids, as well as a number of other lipid metabolites, in the face of normal frequencies of circulating natural killer T (NKT) cells. Specific comparison of FA and asthmatic subjects revealed differences in the microbiota-sensitive aromatic amino acid and secondary bile acid metabolism. Children with both FA and asthma exhibited a metabolomic profile that aligned with that of FA alone but not asthma. Among children with FA, distinct profiles were associated with history of severe reactions and presence of multiple FA.

**Conclusions:** Children with FA display a disease-specific metabolomic profile that is informative of disease mechanisms and severity, and which dominates in the presence of asthma. Lower levels of sphingolipids and ceramides and other metabolomic alterations observed in FA children may reflect the interplay between an altered microbiota and immune cell subsets in the gut.

## Introduction

Allergic diseases, including food allergy (FA) and asthma, exert significant morbidity worldwide, with FA affecting up to 8% of children and 4-10% of adults in the US alone [1-4]. Yet, the availability of accurate biomarkers and effective treatments lags behind the ongoing FA epidemic, leaving providers with imperfect diagnostic tools and patients with food avoidance as their main treatment option. Allergic disorders are complex conditions that result from the interaction of genetic, epigenetic and environmental influences [5]. Though allergic disorders are defined by their shared type 2 immune mechanisms [6], their markedly diverse clinical manifestations suggest that additional unique pathogenic processes are involved in shaping their heterogeneous phenotypic attributes. A deeper understanding of such pathogenic mechanisms may lead to much needed improved disease biomarkers of disease as well as to novel treatment strategies.

Untargeted metabolomic profiling has emerged as a powerful technique to dissect altered pathways contributing to complex diseases [7-10]. Its ability to detect alterations that are influenced by genetic and/or environmental factors makes it particularly useful for unveiling disease mechanisms and for biomarker discovery in complex atopic disorders. Several studies have applied untargeted metabolomics to the study of asthma and have identified metabolic signatures associated with severity and endotypes of asthma [11-13]. In comparison, very few studies have applied untargeted metabolomics to investigate FA. One murine study showed alterations in uric acid metabolism associated with peanut allergy, leading the authors to hypothesize a role for alarmins in FA pathogenesis [14]. More recently, a study of fecal metabolomics suggested that intestinal *Bacteroides*-derived sphingolipids, and particularly *B fragilis* derived alpha-galactosylceramide, were associated with protection against clinical FA in food-sensitized children, possibly through modulation of iNKT cells function [15].

In this report, we performed untargeted metabolomic profiling in FA children with and without asthma, in children with asthma and in non-atopic controls, to better define metabolic changes associated with each clinical entity. These studies aimed to identify metabolites that uniquely correlate with the presence of FA and to gain insight into underlying mechanisms involved in disease pathogenesis. Results revealed unique metabolomic signatures associated with FA that shed light on underlying pathophysiological mechanisms.

## Methods

### Study Population

A total of 125 subjects were included in this study: 35 children with FA alone, 35 with asthma alone, 35 with both FA and asthma and 20 non-atopic controls. Children ages 12 and younger were included, with only non-atopic controls including subjects up to 18 years of age. Participants were recruited in the Allergy clinic at Boston Children’s Hospital and from the SICAS (School Inner City Asthma Study), a NIH funded prospective study of environmental risk factors in inner city school children with asthma (PI, Dr Phipatanakul) [16]. The study was approved by the Boston Children’s Hospital IRB.

FA diagnosis was made by the treating physician and was based on positive IgE testing (skin testing and/or specific IgE) coupled with recent history of IgE-mediated symptoms (hives, angioedema, emesis, diarrhea, wheezing, hypotension etc.) developing within 2 hours of ingestion of the culprit food. Asthma diagnosis was based on validated questionnaire and ATS criteria [16]. Asthmatic patients requiring treatment with systemic steroids were excluded. Non-atopic controls had no history of IgE mediated conditions and/or negative IgE-based testing. Exclusion criteria included history of non-IgE mediated or chronic inflammatory gastrointestinal disorders, use of immunosuppressive medications, and use of antibiotics within the previous 6 weeks.

Children with FA were further classified based on the number of MD diagnosed FA (one vs. multiple) and the history of MD-reported systemic reactions requiring use of intramuscular epinephrine.

### Sample preparation

Blood samples were collected in serum-separator tubes and serum stored at -80C within 2 hours of collection, according to recommended processing guidelines for metabolomic studies.

### Metabolomic profiling

Non-targeted global metabolomic profiles were generated through Metabolon Inc, (Research Triangle, NC) utilizing ultra-performance liquid chromatography coupled with high resolution/accurate mass spectrometer (UPLC-MS/MS). Four platforms were used to detect a comprehensive list of metabolites: 1) UPLC-MS/MS under positive ionization, 2) UPLC-MS/MS under negative ionization, 3) UPLC-MS/MS polar platform (negative ionization) and 4) gas chromatography-MS. Metabolites were identified by their m/z retention time, and through comparison to library entities of purified known standards.

### iNKT cell analyses

Peripheral blood mononuclear cells (PBMCs) were isolated by Ficoll gradient sedimentation according to standard protocols. NK, NKT and iNKT cells were identified as CD3^-^ CD56^+^, CD3^+^ CD56^+^ and CD3^+^ CD56^+^V24Ja18^+^ respectively (Biolegend antibodies). iNKT cells were expressed as a percentage of PBMCs and compared between subjects with FA, asthma and non-atopic controls. Mann-Whitney test was used to compare iNKT cell proportions and significance level was set at p ≤0.05.

### Statistical Analysis

Pairwise comparisons were performed to identify metabolites and pathways dysregulated among different groups. Following log-transformation and imputation of missing values, if any, with the minimum observed value for each compound, each biochemical was rescaled to set the median equal to 1 (Scaled Value). Welch T test, Mann-Whitney and ANOVA contrasts were used to identify biochemicals that differed significantly between experimental groups. To account for multiple comparisons a stringent level of significance of p<0.005 (equivalent to FDR <0.15 with Benjamini-Hochberg correction) was applied [17]. Pair-wise unadjusted contrasts were also applied to compare groups of FA children with differential severity phenotypes based on number of FA and history of anaphylaxis. Statistical analyses were carried out using Graphpad Prism version 7a and SPSS Statistics version 24.

Random Forest Analysis was performed to identify the most differentially expressed metabolites in pair-wise group comparisons. Using a 50,000-tree forest size, importance score was used to identify the top 30 ranking metabolites. Pathway enrichment analysis - mapping to Metabolon proprietary database - was applied to identify metabolic pathways with metabolites that are over-represented and are therefore targets of significant perturbations under different disease conditions. A pathway enrichment value >1 indicates that the pathway contains more experimentally regulated compounds relative to the study overall. Pathway enrichment: (# of significant metabolites in pathway (k)/ total # of detected metabolites in pathway (m)) / (total # of significant metabolites (n) / total # of detected metabolites (N)), or (k/m)/(n/N).

Regression analyses were performed to adjust for characteristics showing statistically different distributions among study groups (atopic dermatitis and, partially, age). Adjusted analyses resulted in a smaller number of significant metabolites, though all metabolomic pathways detected in unadjusted analyses were confirmed to be dysregulated in adjusted comparisons.

## Results

### Characteristics of study subjects

Population demographics and disease attributes are summarized in **Table 1**. A slightly higher, though not statistically significant, proportion of male participants was present in all groups except for healthy controls. The median age was 4.1 years for children with FA alone, 5.5 years for children with FA and asthma, 7.5 years for children with asthma alone and 6.3 years for controls. A marginally significant (p<0.05) age difference was present between children with FA alone and asthma alone. Differences in race distributions were observed, likely resulting from underlying recruitment strategies, with a higher proportion of non-Caucasian children in the asthma-alone group. A slightly higher rate of allergic rhinitis prevalence was observed in children with asthma – with and without FA - as compared with those with FA alone. Atopic dermatitis was significantly more frequent in children with FA – with or without asthma - as compared to those with asthma alone. Total serum IgE concentrations were available for the majority of children with FA and were significantly higher in those with concomitant asthma. Total IgE concentrations were significantly lower in non-atopic controls. Among FA children, about 30% of individuals had one FA while the remainder had allergies to two or more foods. History of anaphylaxis was similar in FA children with and without asthma.

**Table 1.** Demographic characteristics of the study population.

|  | **Asthma**<br>**(n=35)** | **Food allergy**<br>**(n=35)** | **Food allergy and asthma**<br>**(n=35)** | **Control**<br>**(n=20)** | **P-value** |
| --- | --- | --- | --- | --- | --- |
| Age (year) | 7.5* | 4.1* | 5.5 | 6.3 | *<0.05 |
| Number (Male/Female) | 35<br>(20/15) | 35<br>(19/16) | 35<br>(21/14) | 20<br>(10/10) | n.s. |
| BMI (median percentile for age) | 61 | 66.5 | 60.5 | 70 | n.s. |
| Self-reported Race %<br>• White<br>• African/American<br>• Hispanic<br>• Asian<br>• Other/n.a. | 17<br>40<br>37<br>0<br>6 | 60<br>11<br>9<br>9<br>11 | 46<br>14<br>28<br>6<br>6 | 40<br>20<br>35<br>0<br>5 |  |
| Allergic Rhinitis | 27/35<br>(77%) | 23/35<br>(67%) | 30/35<br>(86%) | 0/35 | n.s. |
| Atopic Dermatitis | 8/35*<br>(23%) | 21/35*<br>(60%) | 20/35*<br>(57%) | 0/35 | *< 0.05 |
| Total IgE (geometric mean, kU/L) | n.a. | 219** | 547** | 26.5** | **<0.005 |
| Number of food allergies (%)<br>• One<br>• Two or more |  | 29<br>71 | 31<br>69 |  |  |
| Anaphylaxis % |  | 23 | 29 |  |  |

### Global metabolomic profiling

A total of 1,165 biochemicals - 868 compounds of known identity (named biochemicals) and 297 compounds of unknown structural identity (unnamed biochemicals) - were detected in this study. After removal of drug- and xeno-metabolites, 733 compounds were included in the analyses. Levels of gonadal steroid metabolites strongly correlated with age. Differences in this category of steroids are not reported as they are considered to reflect age effects rather than true pathophysiological characteristics of the study groups.

### Comparisons of children with FA, asthma and controls

Pair-wise comparison of FA children and non-atopic controls revealed 53 metabolites that differed significantly at a stringent cutoff p value of <0.005 (**Figure 1,** Suppl. Table 1). Prominent among these are lipid metabolites, most notably sphingolipids and ceramides, lysophospholipids, lysoplasmalogens, diacylglycerol and fatty acids. Other metabolites included those of the amino acid lysine, threonine and valine, and the β/γ isoforms of tocopherol. There were no differences in the levels of these 53 metabolites between FA children with and without eczema. In comparison to children with asthma, subjects with FA showed changes in 81 metabolites in pair-wise comparisons, involving the metabolisms of sphingolipids, lysophospholipids, mono- and diacylglycerol, fatty acids (monohydroxy, acyl choline), amino acids (tyrosine, tryptophan, histidine), bile acids and tocopherol. (**Figure 2,** Suppl. Table 2). Interestingly, no differences in the levels of these 81 metabolites was present when comparing children with FA either with or without asthma, suggesting that children with these overlapping allergic phenotypes acquire a metabolomic signature concordant with that of FA alone rather than asthma alone. Nine metabolites were uniquely dysregulated in FA children in comparison to both controls and asthmatic children, including the lysoplasmalogen metabolite 1-(1-enyl-palmitoyl)-GPC, lysophospholipids (1-arachidonoyl-GPC, 1-linoleoyl-GPC, 1-palmitoleoyl-GPC, 1-palmitoyl-GPC), N2-acetyllysine, β/γ tocopherol, and sphingomyelins. Alterations uniquely detected in the comparison of children with FA and asthma alone included changes in secondary bile acids and aromatic amino acid (histidine, tyrosine, tryptophan) metabolisms.

**Figure 1.**
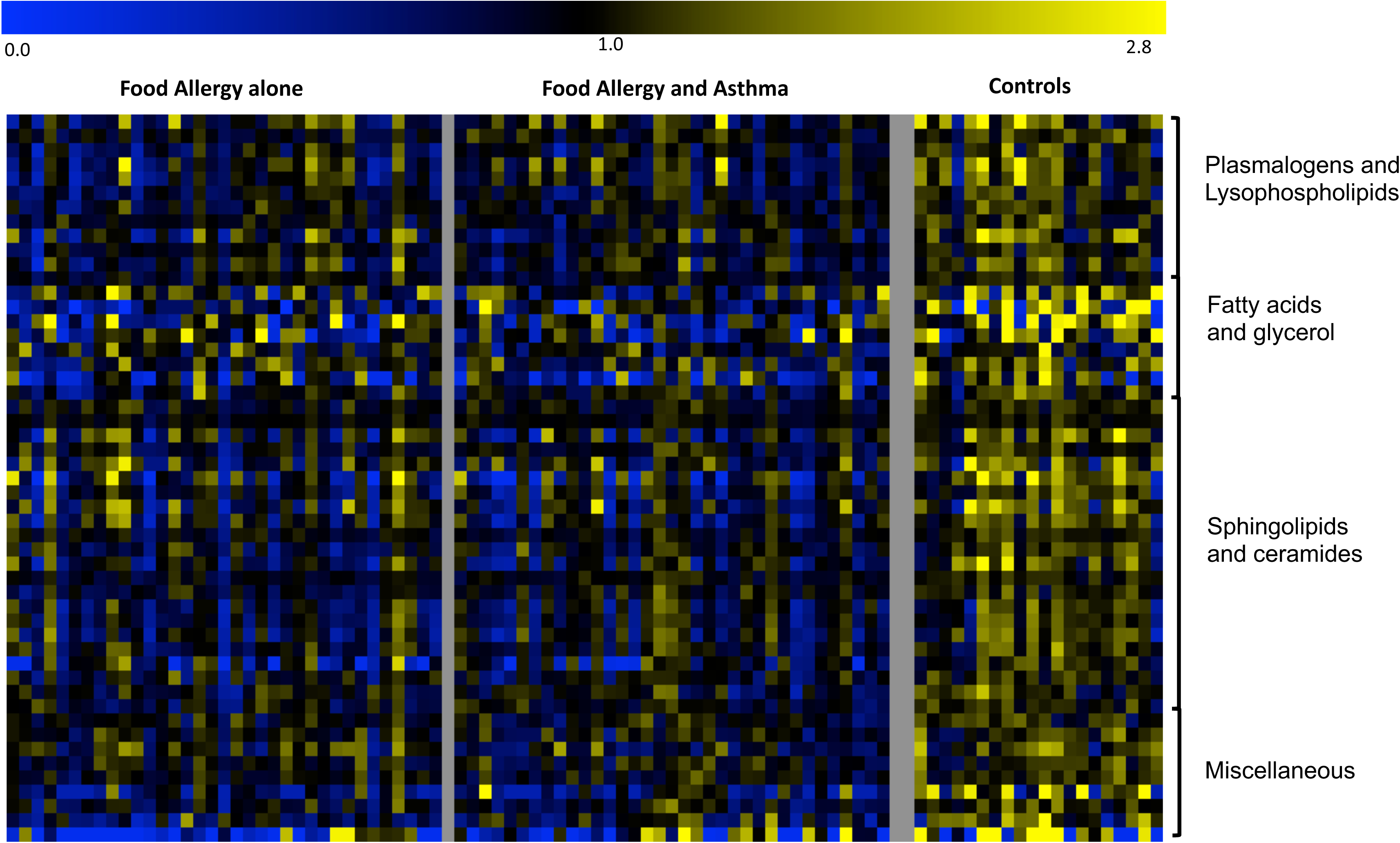
Heat-map of metabolites significantly different (p<0.005) between FA children and non-atopic controls. For each metabolite, a colorimetric representation of relative expression in each sample is shown according to the scale depicted on top. Metabolites are grouped into main dysregulated pathways and a miscellaneous category.

**Figure 2.**
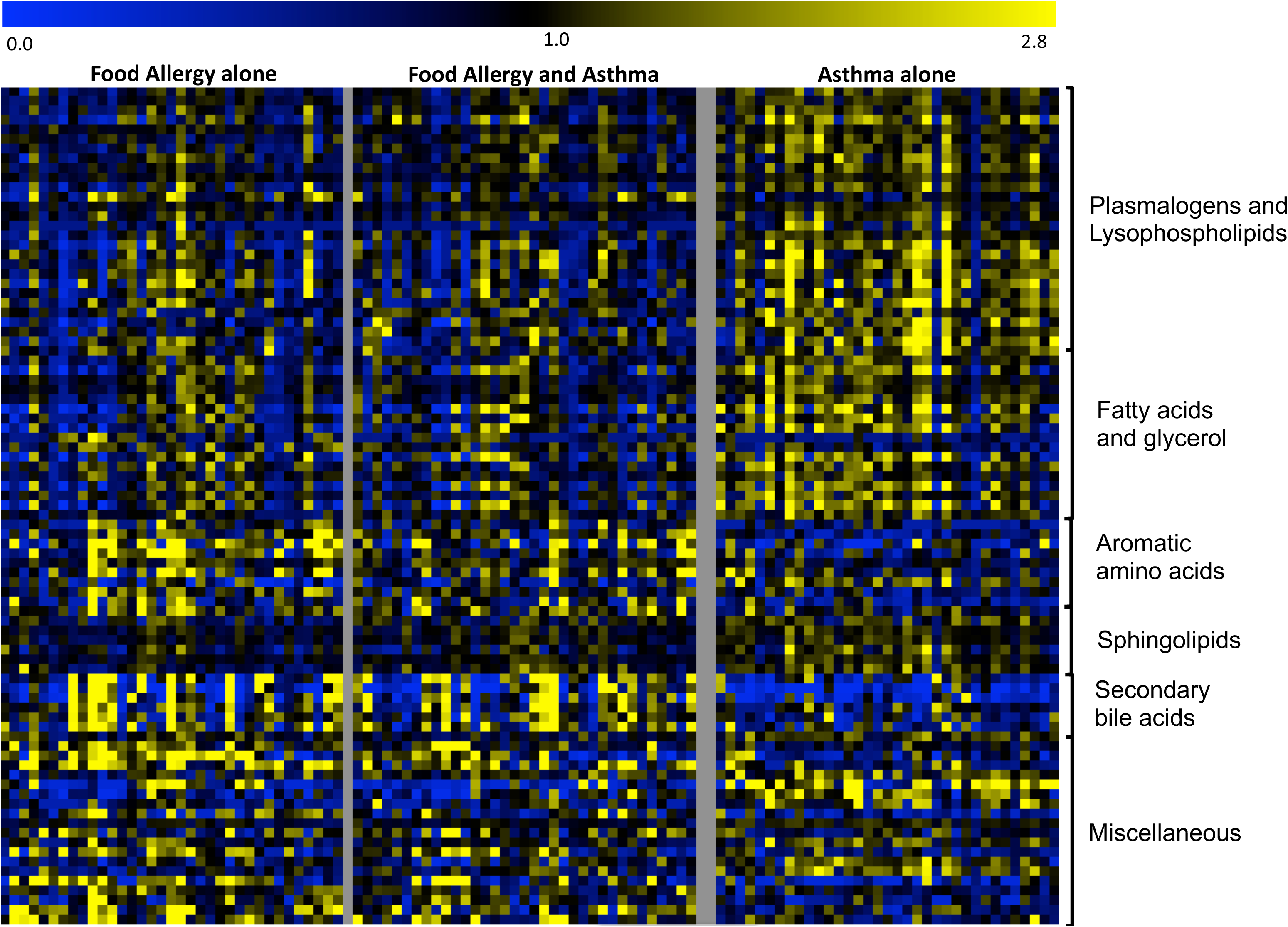
Heat-map of metabolites significantly different (p<0.005) between FA and asthmatic children. For each metabolite, a colorimetric representation of relative expression in each sample is shown according to the scale depicted on top. Metabolites are grouped into main dysregulated pathways and a miscellaneous category.

### Comparisons of children with asthma and controls

Pair-wise contrasts of children with asthma versus non-atopic controls identified 51 metabolites that differed significantly (p<0.005) between the two groups (**Suppl. Figure 1**, Suppl. Table 3) with changes affecting metabolism of plasmalogens, phospholipids, fatty acids (monohydroxy, medium chain, dicarboxylate), sphingolipids, and amino acids (lysine, leucine, valine, threonine). Among these are metabolites previously reported to being associated with asthma phenotypes and pathogenesis, including amino acids (threonine [18-20], valine [21], isoleucine [21], glutamate [22]), primary bile acids (taurocholate) [23], fatty acids (caproate) [24], phosphatidylcholine [25].

### A highly pronounced sphingolipid signature in FA

A particularly prominent pathway affected in FA is sphingolipid metabolism. The steps involved in ceramides and sphingolipid metabolism are schematically shown in **Figure 3**. Key to this pathway is the synthesis of ceramides, and their further metabolisms to sphingomyelins and conjugated ceramides. Several metabolites along this pathway were decreased in FA children as compared to controls and to asthmatics (**Figure 3**). Most of the changes detected in FA subjects appeared to center on the conversion of ceramides to sphingomyelin and acyl- and glucosylceramide, which were decreased in FA (**Fig. 3 A,B**). Alterations in this pathway were also associated with characteristics of FA severity, as described below (**Fig. 3 C,D**).

**Figure 3.**
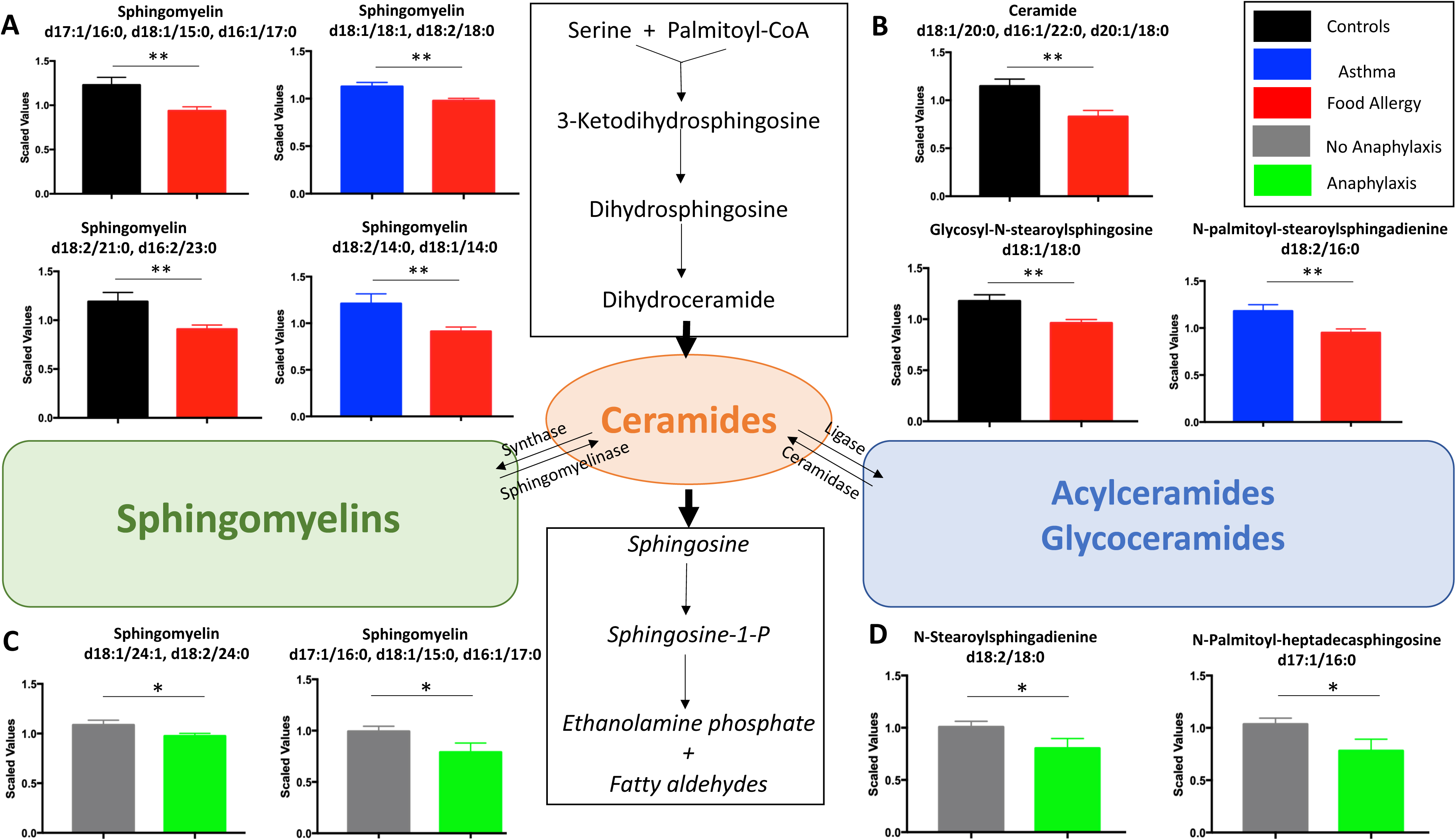
Sphingolipid alterations are prominent in FA. Pairwise comparison of sphingomyelin (**3 A,C**) and ceramide (**3 B,D**) metabolites between children with FA and either asthmatic or non-atopic children as well between FA children with and without asthma are shown. Mean and SEM are shown. * p≤0.05, ** p≤0.005

### FA is not associated with alterations in circulating iNKT cells

iNKT cells represent a small population of circulating NKT cells that carry an invariant T-cell receptor that recognizes lipid moieties, including sphingolipids produced by gut microbial species such as Bacteroidetes, and B. fragilis in particular [26, 27]. A decrease in peripheral iNKT cells has been previously reported in association with FA [28, 29]. We sought to investigate whether changes in iNKT cells are present in children with FA as compared to children with asthma and controls. The proportion of iNKT cells in PBMCs was measured in 10 patients with FA, 12 with asthma, 5 with both FA and asthma and 8 non-atopic controls. No differences in the proportions of NK and NKT cells were detected among the different groups. The proportion of iNKT cells was not different between children with FA and controls though an increased proportion of iNKT was observed in children with asthma, consistent with previous reports [30, 31] (**Figure 4**).

**Figure 4.**
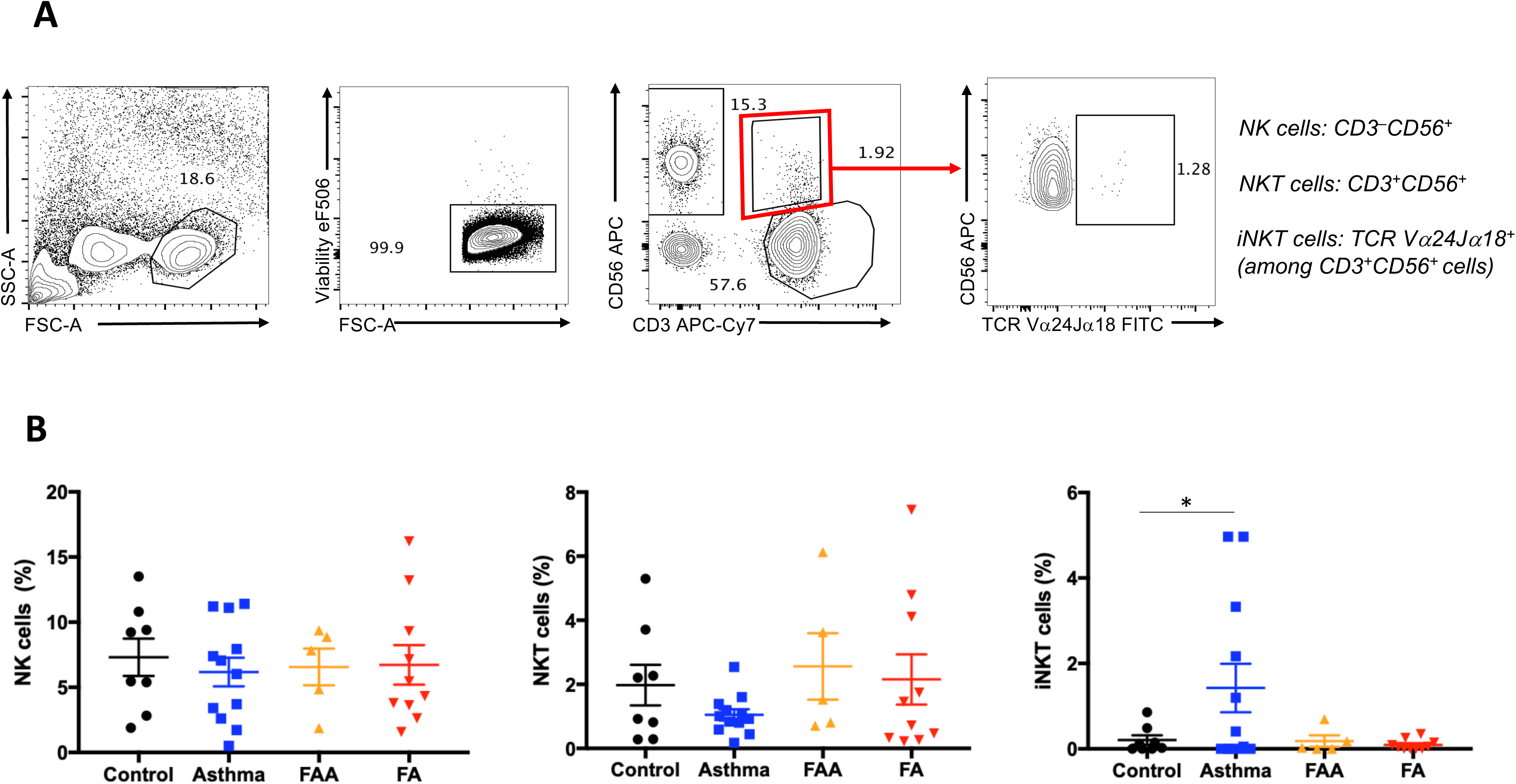
Circulating iNKT cells are unchanged in FA. **4A.** Gating strategy for NK, NKT and iNKT cells. **4B**. Proportions of NK, NKT and iNKT cells in children with FA, asthma, FA and asthma, and non-atopic controls. * p≤0.05

### Metabolomic associations of FA severity phenotypes

FA children were further stratified based on the number of diagnosed FA and on history of systemic reactions requiring IM epinephrine injection as documented by MD.

A total of 41 metabolites in diverse pathways were found to differ between children with FA to one food item (n=21) and children with two or more FA (n=49) (**Suppl. Table 4**). Of interest, children with reactivity to more than one food showed significantly higher levels of plasmalogens and diacylglycerols and lower levels of tryptophan metabolites in the kynurenine and serotonin pathways and of acyl carnitine fatty acids. (**Figure 5, Suppl Table 4**).

**Figure 5.**
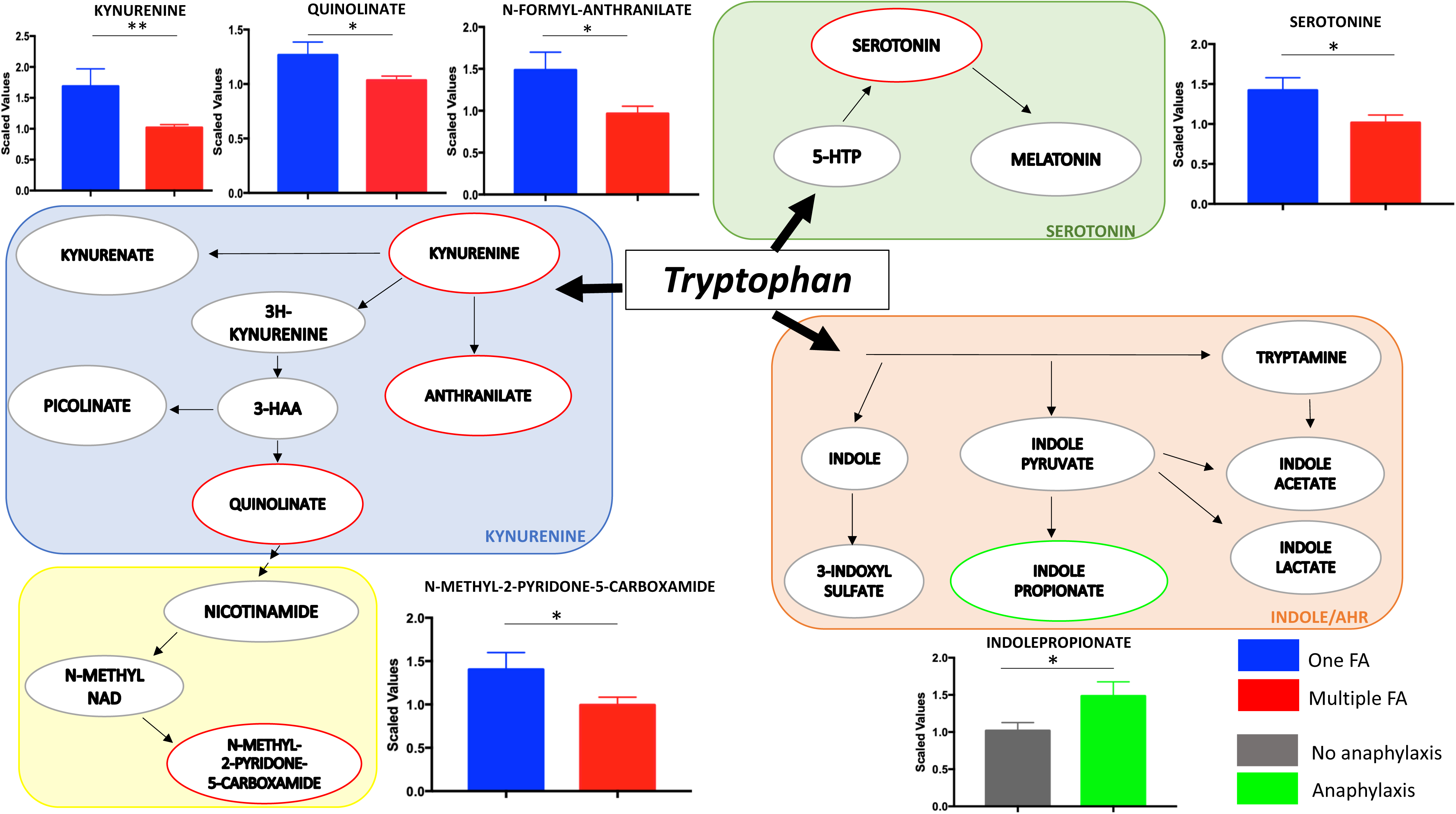
Tryptophan metabolism is dysregulated in children with multiple FA and history of anaphylaxis. The three branches of tryptophan metabolism are depicted together with pairwise comparisons of metabolites significantly different in children with multiple FA or history of anaphylaxis. Mean and SEM are shown. * p≤0.05, ** p≤0.005

A total of 19 metabolites were significantly different when comparing FA children with (n=19) and without (n=51) history of anaphylaxis requiring the use of IM epinephrine. Positive history of anaphylaxis was associated with higher levels of indole tryptophan metabolites and of eicosanoids and lower levels of branched and acyl carnitine fatty acids and sphingolipids (**Figure 5, Suppl. Table 5**).

Interestingly, both characteristics of FA severity were found to be associated with alterations in tryptophan metabolism. Tryptophan is an essential aromatic amino acid that in the gut is under the direct influence of the microbiota and is metabolized through three main pathways depicted in **Figure 5** [32]. While children with more than one FA exhibit a decrease in metabolites of the kynurenine and serotonin pathways, children with positive history of anaphylaxis show a unique increase in the indole pathway as reflected by significantly higher levels of the metabolite indolepropionate.

### Secondary Bile Acid metabolism

Changes in bile acid metabolism were observed in the pairwise comparison of children with FA and asthma (**Figure 2**). Secondary bile acids are derived from primary bile acids by the action of bacterial metabolism and they, in turn, modulate the composition of the gut microbiome [33, 34]. To establish whether the observed differences in bile acid composition attributable to FA may be secondary to altered bacterial metabolism, we compared ratios of different bile acid species that reflect enzymatic conversion steps in the liver and in the gut [35]. A schematic illustration of bile acid metabolism is shown in **Figure 6**. No differences in cholesterol level and metabolism of primary bile acids, including changes between the classic and alternative pathways, were observed between asthmatic children with or without FA. When comparing ratios of microbiome-derived secondary bile acids to liver-generated primary bile acids, children with FA showed increased production of the alternative pathway secondary bile acids ursodeoxycholic acid (UDCA:CDCA ratio), glycoursodeoxycholic acid (GUDCA:CDCA ratio), tauroursodeoxycholic acid (TUDCA:CDCA ratio), and hyocholic acid (HCA:GLCA ratio), and decreased production of glycolithocholic acid (GLCA:CDCA). When testing enzymatic activities related to the conjugation of bile acids to glycine or taurine, the presence of FA was associated with decreased glycine conjugation in secondary bile acids of the classic pathway (GDCA:DCA ratio) and increased taurine conjugation in the alternative pathway (THCA:HCA ratio), suggesting alterations in the processing of primary bile acids by the microbiota.

**Figure 6.**
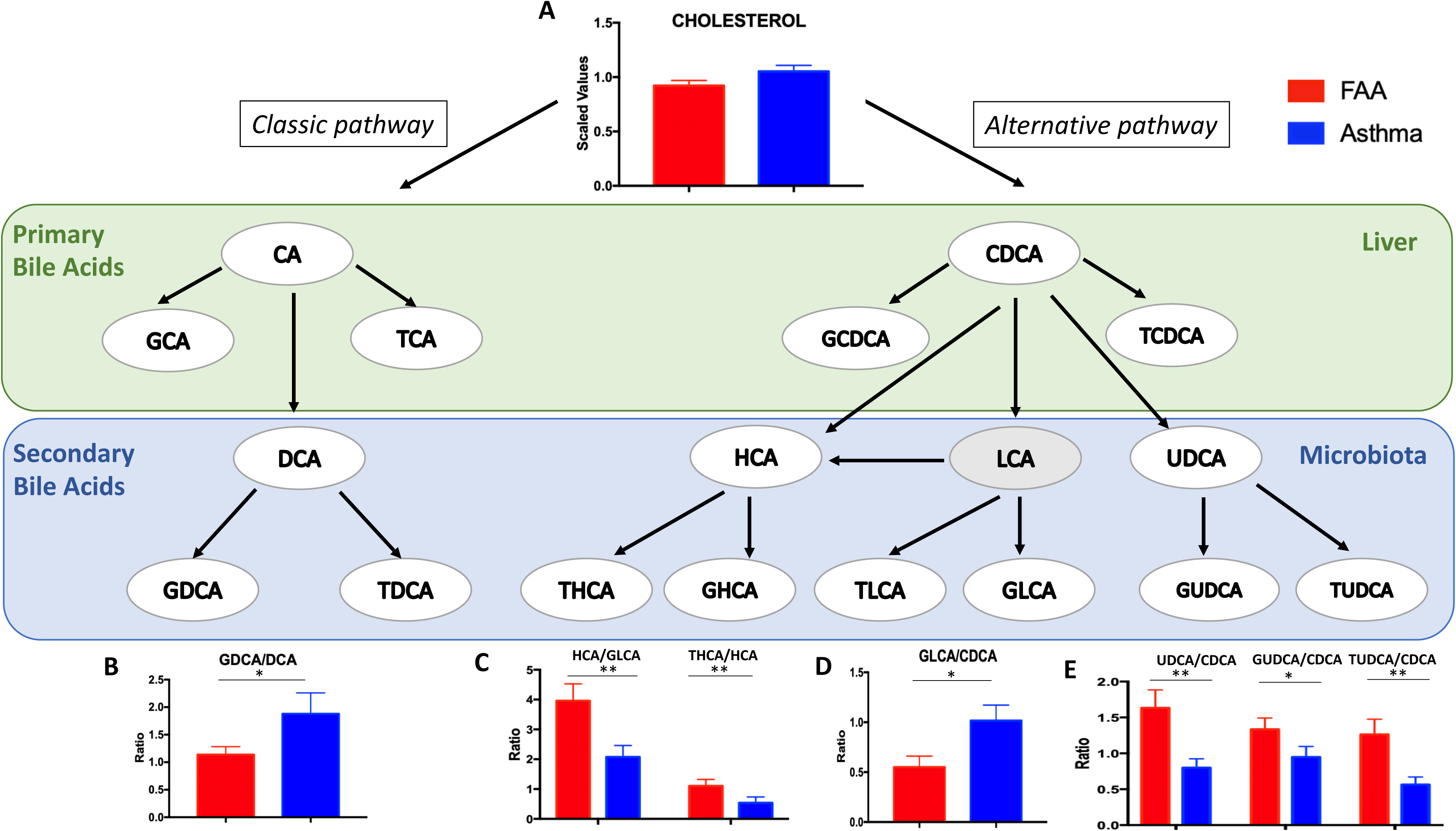
Differences in bile acid ratios between children with FA and asthma. A schematic representation of bile acid metabolism in the liver and the gut is depicted, followed by pairwise comparison of differentially altered bile acid ratios between asthmatic and food allergic children. Mean and SEM are shown. * p≤0.05, ** p≤0.005

### Pathway Analyses

As shown in **Suppl. Figure 2A**, the most dysregulated pathways associated with the presence of FA in comparison to controls include dihydrosphingomyelins, lactosylceramides, lysoplasmalogen, oxidative phosphorylation, sphingomyelins, and hexosylceramides. When compared to asthma (**Suppl. Figure 3A**), the most dysregulated pathways associated with FA included creatine metabolism, fructose/mannose/galactose, mevalonate metabolism, pantothenate and CoA metabolism.

### Random Forest Analysis

Random forest binning of FA vs control samples showed moderate accuracy (70%). Binning of asthma vs control samples and of FA vs asthma samples resulted in overall predictive accuracies of 79% and 87%, significantly better than the 50% that would occur by chance alone. Among FA children, Random forest binning of samples with and without asthma was not predictive, suggesting strong similarity between children with FA alone and those with overlapping FA and asthma. The biochemical importance plot - which displays the top 30 differentiating biochemicals metabolites - in the comparison between FA and controls (**Suppl. Figure 2B**) suggests key differences in membrane lipids (glycerophosphoinositol and sphingomyelin being the most discriminating), vitamins (gamma and beta tocopherol) and amino acids (glutamine, lysine, proline). 1-arachidonoyl-GPC – a plasmalogen and precursor in the metabolism of platelet activating factor - is the highest predictive metabolite of FA versus control. In the comparison of children with FA alone and those with asthma alone (**Suppl. Figure 3B**), the most discriminating metabolites are represented by vitamins/cofactors (gamma- and beta-tocopherol, pantothenate), lipids (glycerophosphoinositol), and aminoacids/peptides (leucylalanine, cysteine-sulfate).

## Discussion

In this study, untargeted serum metabolomic profiling of children with distinct atopic phenotypes led to identification of a unique metabolomic signature associated with FA. This signature was dominant in subjects with both FA and asthma, likely reflecting unique pathogenic mechanisms active in FA that are different from those involved in asthma and, potentially, other atopic disorders.

A hallmark of the FA metabolomic signature is the presence of distinct alterations in lipid metabolites, most prominently a marked decrease in sphingolipids, including sphingomyelins and ceramides. Sphingolipids are a major class of lipids in eukaryotes [36]. *De novo* synthetized sphingolipids are major components of cellular membranes and participate in a multitude of cellular functions [36]. Sphingolipids are also derived from dietary sources and can be produced by *Bacteroides* species in the gut [37]. In a recent study, higher levels of stool sphingolipids were detected in food-sensitized children as compared to subjects with clinical FA or to controls [15]. iNKT cells recognize lipid ligands presented by the atypical major histocompatibility class I molecule CD1d, including *Bacteroides*-derived sphingolipids. In the same study, in-vitro iNKT cell activation by fecal sphingolipids was increased in children with food sensitization but not those with FA. A role for iNKT cells in FA has also been suggested by reports of decreased iNKT cell number in cow milk-allergic children [28] and their increase following milk desensitization [29]. Nevertheless, we did not find differences in the frequency of iNKT cells in FA children versus non-atopic controls, while a small, significant increase in the proportion of iNKT cells was observed in asthmatic children, consistent with previous observations [30, 31]. Sphingolipid changes observed in FA suggest decreased generation of ceramidesand also attenuation of additional distal steps in the conversion of ceramides into sphingomyelins, possibly at the level of the enzymes sphingomyelin synthase and sphingomyelinase. This step in sphingomyelin metabolism has been noted to control the differentiation of Th17 cells [38], suggesting an immune regulatory role for this pathway in modulating regulatory T (Treg) cell/Th17 cell differentiation relevant to FA [39]. Overall, our studies suggest an unexpected and potentially important role for sphingolipid metabolism in FA, and also the potential use of sphingolipid metabolites as disease biomarkers.

FA subjects manifested reductions in other classes of lipids, including lysophospholipids and fatty acids such as mono- and diacylglycerols. We have recently observed similar reduction in lysophospholipids, and especially in levels of plasmalogens, in a mouse model of FA (data not shown). These observations strongly suggest a key role for lipid alterations in the development of FA that is conserved across species and warrants further investigations. Dysregulation in metabolism of selected amino acids (lysine, valine, threonine) and Vitamin E was also associated with FA.

Changes in the microbiome have been linked to allergic disorders, including FA [40] and asthma [41]. We have recently shown a critical role for dysbiosis in the pathogenesis of FA in both mice and humans [39]. Microbiota can affect the host with various mechanisms, including through the production of secreted metabolites [42]. In our study children with FA and asthma displayed significant differences in the ratios of secondary bile acids, which are generated in the gut by the action of anaerobic bacteria. Specifically, we observed increased production of the alternative bile acid UDCA and its conjugates, as well as differences in the rate of taurine/glycine conjugation between the classical and alternative pathways. Our findings not only support a role for dysbiosis in FA and asthma but also suggest that differential signatures of microbial-induced metabolites can be detected in distinct allergic phenotypes and serve as markers of disease. Furthermore, secondary bile acids activate bile acid receptors expressed both in the intestine and other tissues to modulate local and systemic responses, including inflammatory and innate immune responses [43]. Changes in bile acids have been implicated in the pathogenesis of inflammatory disorders [35, 44] and long-ranging effects of dysregulated secondary bile acid metabolism may be present in FA children.

When comparing children with FA and asthma, we observed that children with both conditions have a metabolomic profile that was overlapping with that of children with FA alone. This observation suggests that the presence of FA imparts a metabolomic signature reflective of pathogenic mechanisms in FA that dominate in the co-presence of asthma. Also consistent with distinct disease mechanisms are reports that FA is an independent risk factor for worse asthma outcomes, including frequency and severity of symptoms [45, 46]. These results argue for synergistic interactions between mechanisms operative in different atopic disorders that can influence disease manifestations.

Patients with FA may exhibit severe phenotypes characterized by multiple FA and/or life-threatening systemic reactions. The presence of multiple FA and of a positive history of anaphylaxis were associated with specific metabolomic alterations, including those that contribute to oral tolerance, most prominently the essential aromatic amino acid tryptophan and short chain fatty acids. Tryptophan is metabolized through three different pathways: 1) direct transformation by the gut microbiota into indole metabolites, which can activate the aryl hydrocarbon receptor (AhR); 2) the kynurenine pathway in both immune and epithelial cells by the action of 2,3-dioxygenase-1; and 3) the production of serotonin in the enterochromaffin cells via Trp hydroxylase-1. We found decreased levels of metabolites in the kynurenine and serotonin pathways in children with multiple FA, and a significant increase in the AhR ligand indole-propionate in children with history of anaphylaxis. Both the kynurenine and indole pathways are important in immune responses at mucosal barriers, including Treg cells, and contribute to intestinal homeostasis. Similarly, short chain fatty acids impact Treg cell homing and stability at the barrier interface. Our findings thus indicate that abnormalities in tolerogenic metabolic pathways may be a determinant of disease severity in FA.

A key strength of our study is the recruitment of carefully phenotyped patients with either one or overlapping atopic diseases (FA, Asthma, FAA). MD diagnosis of FA was used, based on the combination of recent occurrence of IgE mediated symptoms and positive IgE-based testing to the culprit food. This approach eliminates known biases related to self-reporting of FA symptoms, which have been shown to not adequately correlate with a physician diagnosis of FA [47]. Another strength is availability of information on disease severity, which allowed for the first-time correlation between the severity of FA and changes in specific metabolomic pathways.

There are also limitations in our study which stem from its pilot nature. Sample sizes were relatively limited. We have addressed this limitation by applying a more stringent significance level (p≤0.005) to report differences among groups. Future studies with larger sample sizes will be needed to validate our findings as well as to perform subgroup analyses. FA children have restricted diets, which may in turn affect their metabolomic profiling. To address this concern, children with different FA were included to counteract the effects of individual food avoidance. While it is possible that dietary practices may still have impacted the metabolomic alterations observed in FA children, many of the changes detected in our study involved metabolites that are not directly related to dietary intake or are contained in a variety of foods that were not differentially avoided by FA subjects. Future studies looking at larger groups of children with individual FA as well as careful dietary recall will be needed to determine the extent of dietary restrictions on the metabolomic profile in FA children.

In summary, our studies established the presence of unique metabolomic signatures associated with FA and disease severity. They also revealed the alignment of the metabolomic signatures of children with FA with and without asthma, indicative of distinct disease mechanisms operative in FA regardless of the presence asthma. Our observations suggest that metabolomic profiling can be a powerful approach in advancing the knowledge into FA pathogenesis and a useful tool in identifying candidate biomarkers and targets for therapeutic intervention.

---

**Supplementary Figure 1.** Heat-map of metabolites significantly different (p<0.005) between asthmatic children and non-atopic controls. For each metabolite, a colorimetric representation of relative expression in each sample is shown according to the scale depicted on top. Metabolites are grouped into main dysregulated pathways and a miscellaneous category.

**Supplementary Figure 2.** Pathway analysis and Random Forrest Analysis of metabolites significantly dysregulated between FA children and controls. Panel **A** shows the fold increase over controls of the most dysregulated pathways in children with FA. Panel **B** shows the top thirty metabolites that most strongly discriminate between children with FA and controls.

**Supplementary Figure 3.** Pathway analysis and Random Forrest Analysis of metabolites significantly dysregulated between FA and asthmatic children. Panel **A** shows the fold increase over controls of the most dysregulated pathways in children with FA. Panel **B** shows the top thirty metabolites that most strongly discriminate between children with FA and asthma.

**Supplementary Table 1**. List of metabolites significantly different between FA children and controls. Metabolite name, pathway, and p-values are listed.

**Supplementary Table 2**. List of metabolites significantly different between FA and asthmatic children. Metabolite name, pathway, and p-values are listed.

**Supplementary Table 3**. List of metabolites significantly different between children with asthma and controls. Metabolite name, pathway, and p-values are listed.

**Supplementary Table 4.** List of metabolites significantly different between FA children with allergy to either one or multiple food categories. Metabolite name, pathway, and p-values are listed.

**Supplementary Table 5.** List of metabolites significantly different between FA children with and without history of anaphylaxis. Metabolite name, pathway, and p-values are listed.

