## Supplemental tables for "Untargeted Metabolomic Profiling Identifies Disease-specific Profiles in Food Allergy"

**Supplementary Table 1**. List of metabolites statistically significant ($p \leq 0.005$) in pairwise comparisons between children with FA and non-atopic controls. Metabolite name, pathway, and p-values are listed.

| | Metabolite | Pathway | P-value |
|---|---|---|---|
| **1** | 1-(1-enyl-palmitoyl)-2-palmitoyl-GPC (P-16:0/16:0) | Plasmalogen | 0.002 |
| **2** | 1-(1-enyl-oleoyl)-GPE (P-18:1) | Lysoplasmalogen | 0.002 |
| **3** | 1-(1-enyl-stearoyl)-GPE (P-18:0) | Lysoplasmalogen | 0.0004 |
| **4** | 1-(1-enyl-palmitoyl)-GPC (P-16:0) | Lysophospholipid | 0.001 |
| **5** | 1-(1-enyl-palmitoyl)-GPE (P-16:0) | Lysophospholipid | 0.005 |
| **6** | 1-arachidonoyl-GPC (20:4n6) | Lysophospholipid | 0.00003 |
| **7** | 1-arachidonoyl-GPE (20:4n6) | Lysophospholipid | 0.001 |
| **8** | 1-linolenoyl-GPC (18:3) | Lysophospholipid | 0.005 |
| **9** | 1-linoleoyl-GPC (18:2) | Lysophospholipid | 0.001 |
| **10** | 1-oleoyl-GPC (18:1) | Lysophospholipid | 0.003 |
| **11** | 1-palmitoleoyl-GPC (16:1) | Lysophospholipid | 0.002 |
| **12** | 1-palmitoyl-GPC (16:0) | Lysophospholipid | 0.003 |
| **13** | glycerophosphoinositol | Phospholipid | 0.00002 |
| **14** | 1-linoleoyl-2-arachidonoyl-GPC (18:2/20:4n6) | Phosphatidylcholine (PC) | 0.004 |
| **15** | 1-myristoyl-2-arachidonoyl-GPC (14:0/20:4) | Phosphatidylcholine (PC) | 0.004 |
| **16** | 3-hydroxyhexanoate | Fatty Acid, Monohydroxy | 0.004 |
| **17** | heptenedioate (C7:1-DC) | Fatty Acid, Dicarboxylate | 0.004 |
| **18** | octadecanedioate (C18-DC) | Fatty Acid, Dicarboxylate | 0.002 |
| **19** | stearoylcarnitine (C18) | Fatty Acid Metabolism (Acyl Carnitine) | 0.003 |
| **20** | oleoyl-arachidonoyl-glycerol (18:1/20:4) | Diacylglycerol | 0.001 |
| **21** | palmitoyl-arachidonoyl-glycerol (16:0/20:4) | Diacylglycerol | 0.004 |

| | | | |
|---|---|---|---|
| **22** | stearoyl-arachidonoyl-glycerol (18:0/20:4) | Diacylglycerol | 0.002 |
| **23** | ceramide (d18:1/20:0, d16:1/22:0, d20:1/18:0) | Ceramides | 0.002 |
| **24** | glycosyl-N-stearoyl-sphingosine (d18:1/18:0) | Ceramides | 0.003 |
| **25** | lactosyl-N-nervonoyl-sphingosine (d18:1/24:1) | Ceramides | 0.001 |
| **26** | lactosyl-N-palmitoyl-sphingosine (d18:1/16:0) | Ceramides | 0.001 |
| **27** | behenoyl dihydrosphingomyelin (d18:0/22:0) | Sphingolipid Metabolism | 0.004 |
| **28** | lignoceroyl sphingomyelin (d18:1/24:0) | Sphingolipid Metabolism | 0.003 |
| **29** | myristoyl dihydrosphingomyelin (d18:0/14:0) | Sphingolipid Metabolism | 0.001 |
| **30** | palmitoyl dihydrosphingomyelin (d18:0/16:0) | Sphingolipid Metabolism | 0.005 |
| **31** | palmitoyl sphingomyelin (d18:1/16:0) | Sphingolipid Metabolism | 0.004 |
| **32** | sphingomyelin (d17:1/14:0, d16:1/15:0) | Sphingolipid Metabolism | 0.002 |
| **33** | sphingomyelin (d17:1/16:0, d18:1/15:0, d16:1/17:0) | Sphingolipid Metabolism | 0.003 |
| **34** | sphingomyelin (d18:0/20:0, d16:0/22:0) | Sphingolipid Metabolism | 0.0002 |
| **35** | sphingomyelin (d18:1/14:0, d16:1/16:0) | Sphingolipid Metabolism | 0.001 |
| **36** | sphingomyelin (d18:1/22:1, d18:2/22:0, d16:1/24:1) | Sphingolipid Metabolism | 0.001 |
| **37** | sphingomyelin (d18:1/22:2, d18:2/22:1, d16:1/24:2) | Sphingolipid Metabolism | 0.0005 |
| **38** | sphingomyelin (d18:2/14:0, d18:1/14:1) | Sphingolipid Metabolism | 0.002 |
| **39** | sphingomyelin (d18:2/16:0, d18:1/16:1) | Sphingolipid Metabolism | 0.00001 |
| **40** | sphingomyelin (d18:2/18:1) | Sphingolipid Metabolism | 0.001 |
| **41** | sphingomyelin (d18:2/21:0, d16:2/23:0) | Sphingolipid Metabolism | 0.004 |
| **42** | sphingomyelin (d18:2/23:0, d18:1/23:1, d17:1/24:1) | Sphingolipid Metabolism | 0.001 |
| **43** | sphingomyelin (d18:2/23:1) | Sphingolipid Metabolism | 0.001 |
| **44** | tricosanoyl sphingomyelin (d18:1/23:0) | Sphingolipid Metabolism | 0.002 |

| | | | |
|---|---|---|---|
| **45** | taurodeoxycholate | Secondary Bile Acid Metabolism | 0.0004 |
| **46** | lysine | Lysine Metabolism | 0.0001 |
| **47** | N2-acetyllysine | Lysine Metabolism | 0.0005 |
| **48** | threonine | Glycine, Serine and Threonine Metabolism | 0.002 |
| **49** | valine | Leucine, Isoleucine and Valine Metabolism | 0.003 |
| **50** | isoleucine | Leucine, Isoleucine and Valine Metabolism | 0.005 |
| **51** | glutamate | Glutamate Metabolism | 0.004 |
| **52** | cholesterol | Sterol | 0.0002 |
| **53** | gamma-tocopherol/beta-tocopherol | Tocopherol Metabolism | 0.003 |

**Supplementary Table 2**. List of metabolites statistically significant ($p \leq 0.005$) in pairwise comparisons between children with FA and non-atopic controls. Metabolite name, pathway, and p-values are listed.

| | Metabolite | Pathway | P-value |
|---|---|---|---|
| **1** | 1-(1-enyl-palmitoyl)-GPC (P-16:0) | Lysoplasmalogen | 0.00003 |
| **2** | 1-arachidonoyl-GPC (20:4n6) | Lysophospholipid | 0.00004 |
| **3** | 1-arachidonoyl-GPI (20:4) | Lysophospholipid | 0.003 |
| **4** | 1-linoleoyl-GPC (18:2) | Lysophospholipid | 0.0001 |
| **5** | 1-linoleoyl-GPG (18:2) | Lysophospholipid | 0.001 |
| **6** | 1-linoleoyl-GPI (18:2) | Lysophospholipid | 0.00004 |
| **7** | 1-palmitoleoyl-GPC (16:1) | Lysophospholipid | 0.002 |
| **8** | 1-palmitoyl-GPC (16:0) | Lysophospholipid | 0.0002 |
| **9** | 1-palmitoyl-GPI (16:0) | Lysophospholipid | 0.00005 |
| **10** | 1-stearoyl-GPC (18:0) | Lysophospholipid | 0.00002 |
| **11** | 1-stearoyl-GPG (18:0) | Lysophospholipid | 0.002 |
| **12** | 1-palmitoyl-2-linoleoyl-GPI (16:0/18:2) | Phosphatidylinositol (PI) | 0.00002 |
| **13** | glycerophosphoinositol | Phospholipid Metabolism | 0.00002 |
| **14** | glycerol | Glycerolipid Metabolism | 0.002 |
| **15** | 1-arachidonylglycerol (20:4) | Monoacylglycerol | 0.0003 |
| **16** | 1-dihomo-linolenylglycerol (20:3) | Monoacylglycerol | 0.000009 |
| **17** | 1-linolenoylglycerol (18:3) | Monoacylglycerol | 0.0005 |
| **18** | 1-linoleoylglycerol (18:2) | Monoacylglycerol | 0.00006 |
| **19** | 1-oleoylglycerol (18:1) | Monoacylglycerol | 0.005 |
| **20** | 1-palmitoleoylglycerol (16:1) | Monoacylglycerol | 0.002 |
| **21** | 2-linoleoylglycerol (18:2) | Monoacylglycerol | 0.001 |

| | | | |
|---|---|---|---|
| **22** | diacylglycerol (16:1/18:2 [2], 16:0/18:3 [1]) | Diacylglycerol | 0.0002 |
| **23** | palmitoleoyl-linoleoyl-glycerol (16:1/18:2) | Diacylglycerol | 0.003 |
| **24** | palmitoyl-linoleoyl-glycerol (16:0/18:2) | Diacylglycerol | 0.0004 |
| **25** | linoleoyl-arachidonoyl-glycerol (18:2/20:4) | Diacylglycerol | 0.003 |
| **26** | 2-hydroxypalmitate | Fatty Acid, Monohydroxy | 0.003 |
| **27** | 2-hydroxystearate | Fatty Acid, Monohydroxy | 0.0004 |
| **28** | arachidonate (20:4n6) | Polyunsaturated Fatty Acid (n3 and n6) | 0.001 |
| **29** | dihomo-linolenate (20:3n3 or n6) | Polyunsaturated Fatty Acid (n3 and n6) | 0.001 |
| **30** | docosapentaenoate (n6 DPA; 22:5n6) | Polyunsaturated Fatty Acid (n3 and n6) | 0.002 |
| **31** | dihomo-linolenoyl-choline | Fatty Acid Metabolism (Acyl Choline) | 0.004 |
| **32** | arachidonoylcholine | Fatty Acid Metabolism (Acyl Choline) | 0.001 |
| **33** | linoleoylcholine | Fatty Acid Metabolism (Acyl Choline) | 0.001 |
| **34** | nervonoylcarnitine (C24:1) | Fatty Acid Metabolism(Acyl Carnitine) | 0.002 |
| **35** | oleoylcholine | Fatty Acid Metabolism (Acyl Choline) | 0.006 |
| **36** | palmitoylcarnitine (C16) | Fatty Acid Metabolism(Acyl Carnitine) | 0.007 |
| **37** | palmitoylcholine | Fatty Acid Metabolism (Acyl Choline) | 0.002 |
| **38** | stearoylcholine | Fatty Acid Metabolism (Acyl Choline) | 0.001 |
| **39** | malonylcarnitine | Fatty Acid Synthesis | 0.001 |
| **40** | stearate (18:0) | Long Chain Fatty Acid | 0.004 |
| **41** | palmitate (16:0) | Long Chain Fatty Acid | 0.004 |
| **42** | 1-methyl-4-imidazoleacetate | Histidine Metabolism | 0.001 |
| **43** | 1-ribosyl-imidazoleacetate | Histidine Metabolism | 0.0004 |
| **44** | 4-imidazoleacetate | Histidine Metabolism | 0.000003 |
| **45** | N-acetylcarnosine | Histidine Metabolism | 0.000003 |

| | | | |
|---|---|---|---|
| **46** | N-acetylhistidine | Histidine Metabolism | 0.00007 |
| **47** | 3-(4-hydroxyphenyl)lactate | Tyrosine Metabolism | 0.001 |
| **48** | 3-methoxytyramine sulfate | Tyrosine Metabolism | 0.0003 |
| **49** | 4-methoxyphenol sulfate | Tyrosine Metabolism | 0.0003 |
| **50** | phenol glucuronide | Tyrosine metabolism | 0.0002 |
| **51** | phenol sulfate | Tyrosine metabolism | 0.003 |
| **52** | 5-hydroxyindoleacetate | Tryptophan Metabolism | 0.001 |
| **53** | N-palmitoyl-sphingadienine (d18:2/16:0) | Sphingolipid Metabolism | 0.002 |
| **54** | sphingomyelin (d18:1/18:1, d18:2/18:0) | Sphingolipid Metabolism | 0.002 |
| **55** | sphingomyelin (d18:1/20:1, d18:2/20:0) | Sphingolipid Metabolism | 0.001 |
| **56** | sphingomyelin (d18:2/14:0, d18:1/14:1) | Sphingolipid Metabolism | 0.003 |
| **57** | sphingomyelin (d18:2/16:0, d18:1/16:1) | Sphingolipid Metabolism | 0.002 |
| **58** | sphingomyelin (d18:2/24:2) | Sphingolipid Metabolism | 0.002 |
| **59** | glycohyocholate | Secondary Bile Acid Metabolism | 0.002 |
| **60** | glycoursodeoxycholate | Secondary Bile Acid Metabolism | 0.004 |
| **61** | taurocholate | Primary Bile Acid Metabolism | 0.003 |
| **62** | taurohyocholate | Secondary Bile Acid Metabolism | 0.001 |
| **63** | tauroursodeoxycholate | Secondary Bile Acid Metabolism | 0.001 |
| **64** | aspartate | Alanine and Aspartate Metabolism | 0.0001 |
| **65** | beta-citrylglutamate | Glutamate Metabolism | 0.006 |
| **66** | N-acetyl-aspartyl-glutamate (NAAG) | Glutamate Metabolism | 0.0001 |
| **67** | gamma-glutamylalanine | Gamma-glutamyl Amino Acid | 0.001 |
| **68** | N2-acetyllysine | Lysine Metabolism | 0.001 |

| | | | |
|---|---|---|---|
| **69** | 2-methylserine | Glycine, Serine and Threonine Metabolism | 0.001 |
| **70** | S-methylmethionine | Methionine, Cysteine, SAM and Taurine Metabolism | 0.001 |
| **71** | glycylvaline | Dipeptide | 0.001 |
| **72** | leucylalanine | Dipeptide | 0.000009 |
| **73** | N-acetyl-isoputreanine | Polyamine Metabolism | 0.002 |
| **74** | N-acetylneuraminate | Aminosugar Metabolism | 0.005 |
| **75** | N-delta-acetylornithine | Urea cycle; Arginine and Proline Metabolism | 0.0001 |
| **76** | gamma-tocopherol/beta-tocopherol | Tocopherol Metabolism | 0.000003 |
| **77** | pyridoxate | Vitamin B6 Metabolism | 0.002 |
| **78** | creatinine | Creatine Metabolism | 0.000006 |
| **79** | cysteine-glutathione disulfide | Glutathione Metabolism | 0.001 |
| **80** | dihydroorotate | Pyrimidine Metabolism, Orotate containing | 0.00003 |
| **81** | fumarate | TCA Cycle | 0.002 |

**Supplementary Table 3**. List of metabolites statistically significant (p≤0.005) in pairwise comparisons between children with asthma and non-atopic controls. Metabolite name, pathway, and p-values are listed.

| | Metabolite | Pathway | P-value |
|---|---|---|---|
| **1** | 1-(1-enyl-palmitoyl)-2-oleoyl-GPC (P-16:0/18:1) | Plasmalogen | 0.001 |
| **2** | 1-(1-enyl-oleoyl)-GPE (P-18:1) | Lysoplasmalogen | 0.001 |
| **3** | 1-(1-enyl-stearoyl)-GPE (P-18:0) | Lysoplasmalogen | 0.005 |
| **4** | 1-arachidonoyl-GPE (20:4n6) | Lysophospholipid | 0.002 |
| **5** | 1,2-dipalmitoyl-GPC (16:0/16:0) | Phosphatidylcholine (PC) | 0.001 |
| **6** | 1-myristoyl-2-arachidonoyl-GPC (14:0/20:4) | Phosphatidylcholine (PC) | 0.001 |
| **7** | 1-myristoyl-2-palmitoyl-GPC (14:0/16:0) | Phosphatidylcholine (PC) | 0.002 |
| **8** | 1-palmitoyl-2-stearoyl-GPC (16:0/18:0) | Phosphatidylcholine (PC) | 0.004 |
| **9** | 1-stearoyl-2-arachidonoyl-GPI (18:0/20:4) | Phosphatidylinositol (PI) | 0.003 |
| **10** | 3-hydroxyhexanoate | Fatty Acid, Monohydroxy | 0.0002 |
| **11** | caproate (6:0) | Medium Chain Fatty Acid | 0.001 |
| **12** | heptenedioate (C7:1-DC) | Fatty Acid, Dicarboxylate | 0.0004 |
| **13** | cholesterol | Sterol | 0.005 |
| **14** | oleoyl-arachidonoyl-glycerol (18:1/20:4) | Diacylglycerol | 0.005 |
| **15** | behenoyl dihydrosphingomyelin (d18:0/22:0) | Sphingolipid Metabolism | 0.001 |
| **16** | myristoyl dihydrosphingomyelin (d18:0/14:0) | Sphingolipid Metabolism | 0.0005 |
| **17** | palmitoyl dihydrosphingomyelin (d18:0/16:0) | Sphingolipid Metabolism | 0.0005 |
| **18** | sphingomyelin (d17:1/14:0, d16:1/15:0) | Sphingolipid Metabolism | 0.001 |
| **19** | sphingomyelin (d18:0/20:0, d16:0/22:0) | Sphingolipid Metabolism | 0.001 |
| **20** | sphingomyelin (d18:1/14:0, d16:1/16:0) | Sphingolipid Metabolism | 0.0005 |
| **21** | ceramide (d16:1/24:1, d18:1/22:1) | Ceramides | 0.004 |

| | | | |
|---|---|---|---|
| **22** | lactosyl-N-nervonoyl-sphingosine (d18:1/24:1) | Ceramides | 0.002 |
| **23** | taurochenodeoxycholate | Primary Bile Acid Metabolism | 0.001 |
| **24** | taurocholate | Primary Bile Acid Metabolism | 0.0004 |
| **25** | glycocholate | Primary Bile Acid Metabolism | 0.001 |
| **26** | lysine | Lysine Metabolism | 0.00001 |
| **27** | 2-aminoadipate | Lysine Metabolism | 0.001 |
| **28** | N2-acetyllysine | Lysine Metabolism | 0.001 |
| **29** | leucine | Leucine, Isoleucine and Valine Metabolism | 0.002 |
| **30** | N-acetylleucine | Leucine, Isoleucine and Valine Metabolism | 0.005 |
| **31** | isoleucine | Leucine, Isoleucine and Valine Metabolism | 0.002 |
| **32** | valine | Leucine, Isoleucine and Valine Metabolism | 0.001 |
| **33** | N-acetylvaline | Leucine, Isoleucine and Valine Metabolism | 0.0002 |
| **34** | 3-methylglutaconate | Leucine, Isoleucine and Valine Metabolism | 0.00008 |
| **35** | gamma-glutamylalanine | Gamma-glutamyl Amino Acid | 0.004 |
| **36** | gamma-glutamyl-alpha-lysine | Gamma-glutamyl Amino Acid | 0.002 |
| **37** | gamma-glutamylthreonine | Gamma-glutamyl Amino Acid | 0.003 |
| **38** | glutamate | Glutamate Metabolism | 0.0004 |
| **39** | glutamine | Glutamate Metabolism | 0.004 |
| **40** | N-acetylthreonine | Glycine, Serine and Threonine Metabolism | 0.0003 |
| **41** | sarcosine | Glycine, Serine and Threonine Metabolism | 0.004 |
| **42** | threonine | Glycine, Serine and Threonine Metabolism | 0.001 |
| **43** | methionine | Methionine, Cysteine, SAM and Taurine Metabolism | 0.002 |
| **44** | N-formylmethionine | Methionine, Cysteine, SAM and Taurine Metabolism | 0.003 |

| | | | |
|---|---|---|---|
| **45** | N-acetylmethionine | Methionine, Cysteine, SAM and Taurine Metabolism | 0.002 |
| **46** | N-acetylhistidine | Histidine Metabolism | 0.004 |
| **47** | 4-hydroxyphenylpyruvate | Tyrosine Metabolism | 0.002 |
| **48** | orotate | Pyrimidine Metabolism, Orotate containing | 0.001 |
| **49** | 3-aminoisobutyrate | Pyrimidine Metabolism, Thymine containing | 0.001 |
| **50** | dimethylarginine | Urea cycle; Arginine and Proline Metabolism | 0.005 |
| **51** | N-acetylneuraminate | Aminosugar Metabolism | 0.0002 |

**Supplementary Table 4**. List of metabolites statistically significant ($p \leq 0.05$) in pairwise comparisons between children with FA to one versus multiple food items. Metabolite name, pathway, and p-values are listed.

| | Metabolite | Pathway | P-value |
|---|---|---|---|
| **1** | 1-(1-enyl-palmitoyl)-2-oleoyl-GPC (P-16:0/18:1) | Plasmalogen | 0.047 |
| **2** | 1-(1-enyl-palmitoyl)-2-palmitoleoyl-GPC (P-16:0/16:1) | Plasmalogen | 0.020 |
| **3** | 1-(1-enyl-palmitoyl)-2-palmitoyl-GPC (P-16:0/16:0) | Plasmalogen | 0.035 |
| **4** | 1-methyl-4-imidazoleacetate | Histidine metabolism | 0.025 |
| **5** | 1-myristoyl-2-palmitoyl-GPC (14:0/16:0) | Phosphatidylcholine | 0.029 |
| **6** | glycerol | Glycerolipid Metabolism | 0.042 |
| **7** | 1-myristoylglycerol (14:0) | Monoacylglycerol | 0.024 |
| **8** | 1-pentadecanoylglycerol (15:0) | Monoacylglycerol | 0.007 |
| **9** | oleoyl-arachidonoyl-glycerol (18:1/20:4) | Diacylglycerol | 0.029 |
| **10** | linoleoyl-arachidonoyl-glycerol (18:2/20:4) | Diacylglycerol | 0.007 |
| **11** | arachidoylcarnitine (C20) | Fatty Acid Metabolism (Acyl Carnitine) | 0.049 |
| **12** | cis-4-decenoylcarnitine (C10:1) | Fatty Acid Metabolism (Acyl Carnitine) | 0.043 |
| **13** | arachidonoylcarnitine (C20:4) | Fatty Acid Metabolism (Acyl Carnitine) | 0.034 |
| **14** | dihomo-linoleoylcarnitine (C20:2) | Fatty Acid Metabolism (Acyl Carnitine) | 0.045 |
| **15** | linolenoylcarnitine (C18:3) | Fatty Acid Metabolism (Acyl Carnitine) | 0.012 |
| **16** | linoleoylcarnitine (C18:2) | Fatty Acid Metabolism (Acyl Carnitine) | 0.007 |
| **17** | 13-methylmyristate (i15:0) | Fatty acid, branched | 0.027 |
| **18** | N-trimethyl 5-aminovalerate | Lysine Metabolism | 0.012 |
| **19** | 2-aminoadipate | Lysine metabolism | 0.019 |
| **20** | 3-hydroxy-2-ethylpropionate | Leucine, Isoleucine and Valine metabolism | 0.013 |
| **21** | isobutyrylcarnitine (C4) | Leucine, Isoleucine and Valine Metabolism | 0.019 |
| **22** | tiglylcarnitine (C5:1-DC) | Leucine, Isoleucine and Valine Metabolism | 0.009 |
| **23** | 4-hydroxyphenylpyruvate | Tyrosine metabolism | 0.031 |

| | | | |
|---|---|---|---|
| **24** | Kynurenine | Tryptophan Metabolism | 0.001 |
| **25** | Kynurenate | Tryptophan Metabolism | 0.027 |
| **26** | Serotonin | Tryptophan Metabolism | 0.026 |
| **27** | N-formylanthranilic acid | Tryptophan Metabolism | 0.008 |
| **28** | Formiminoglutamate | Histidine Metabolism | 0.015 |
| **29** | quinolinate | Nicotinate and Nicotinamide Metabolism | 0.019 |
| **30** | N1-Methyl-4-pyridone-3-carboxamide | Nicotinate and Nicotinamide Metabolism | 0.032 |
| **31** | phenylpyruvate | Phenylalanine Metabolism | 0.004 |
| **32** | S-methylcysteine sulfoxide | Methionine, Cysteine, SAM and Taurine Metabolism | 0.029 |
| **33** | N1-methyladenosine | Purine Metabolism, Adenine containing | 0.036 |
| **34** | orotate | Pyrimidine Metabolism, Orotate containing | 0.028 |
| **35** | N-acetylproline | Urea cycle; Arginine and Proline Metabolism | 0.045 |
| **36** | N-acetylneuraminate | Aminosugar Metabolism | 0.038 |
| **37** | sphingomyelin (d18:2/16:0, d18:1/16:1) | Sphingolipid Metabolism | 0.003 |
| **38** | lactosyl-N-palmitoyl-sphingosine (d18:1/16:0) | Ceramides | 0.022 |
| **39** | glycochenodeoxycholate | Primary Bile Acid Metabolism | 0.017 |
| **40** | succinylcarnitine (C4-DC) | TCA Cycle | 0.004 |
| **41** | Cortisone | Corticosteroids | 0.022 |

**Supplementary Table 5**. List of metabolites statistically significant (p≤0.05) in pairwise comparisons between food allergic children with and without history of anaphylaxis. Metabolite name, pathway, and p-values are listed.

| | Metabolite | Pathway | P-value |
|---|---|---|---|
| **1** | 12-HETE | Eicosanoid | 0.007 |
| **2** | 1-linoleoyl-GPA (18:2) | Lysophospholipid | 0.050 |
| **3** | 1-linolenoylglycerol (18:3) | Monoacylglycerol | 0.048 |
| **4** | acetylcarnitine (C2) | Fatty Acid Metabolism (Acyl Carnitine) | 0.047 |
| **5** | ximenoylcarnitine (C26:1) | Fatty Acid Metabolism (Acyl Carnitine) | 0.023 |
| **6** | hexanoylglycine | Fatty Acid Metabolism (Acyl Glycine) | 0.013 |
| **7** | 10-undecenoate (11:1n1) | Medium chain FA | 0.038 |
| **8** | Indolepropionate | Tryptophan Metabolism | 0.031 |
| **9** | 5-bromotryptophan | Tryptophan metabolism | 0.041 |
| **10** | methionine sulfone | Methionine, Cysteine, SAM and Taurine Metabolism | 0.029 |
| **11** | N-acetylproline | Urea cycle; Arginine and Proline Metabolism | 0.019 |
| **12** | N-carbamoylalanine | Alanine and Aspartate Metabolism | 0.005 |
| **13** | lactate | Glycolysis, Gluconeogenesis, and Pyruvate Metabolism | 0.012 |
| **14** | N-palmitoyl-heptadecasphingosine (d17:1/16:0) | Sphingolipid Metabolism | 0.029 |
| **15** | sphingomyelin (d17:1/16:0, d18:1/15:0, d16:1/17:0) | Sphingolipid Metabolism | 0.043 |
| **16** | sphingomyelin (d18:1/24:1, d18:2/24:0) | Sphingolipid Metabolism | 0.033 |
| **17** | Pyridoxal | Vitamin B6 Metabolism | 0.008 |
| **18** | pantothenate | Pantothenate and CoA Metabolism | 0.018 |
| **19** | (N(1) + N(8))-acetylspermidine | Polyamine metabolism | 0.044 |
